## Supplementary Figures for "Accurate prediction of gene deletion phenotypes with Flux Cone Learning"

### SUPPLEMENTARY FIGURES AND TABLES

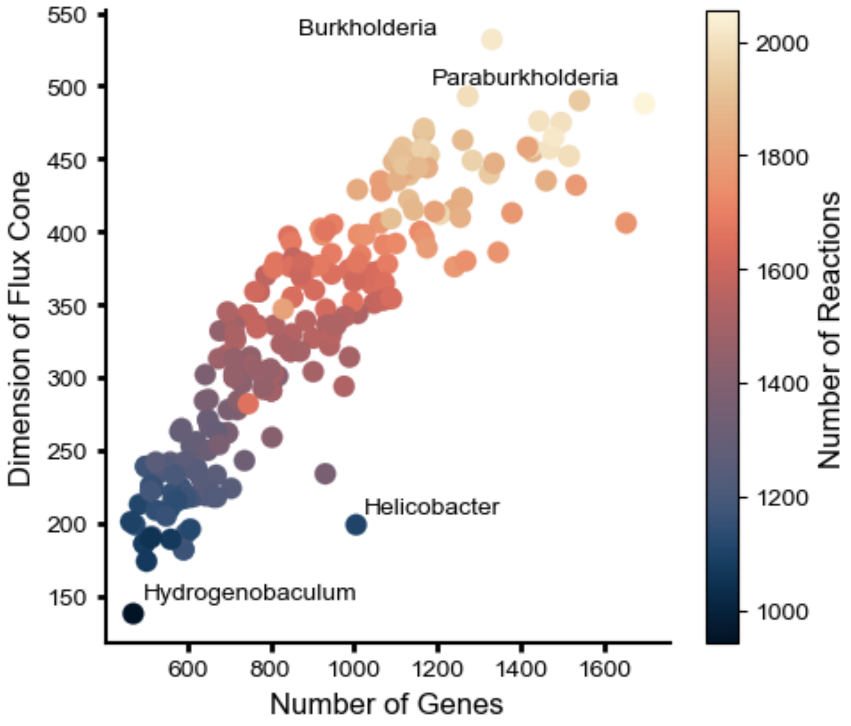


FIG. S1. **Genome-scale metabolic models from across the bacterial kingdom.** Models for $N = 244$ bacterial species were obtained from Plata et al, Nature, 2015 and species names were obtained from Ramon and Stelling, Nature Communications, 2023. The stoichometric matrices of each model were extracted, along with the number of genes (x-axis) and reactions (hue) in each model. The nullity of the stoichiometric matrix (the number of columns in the null space) was computed for each species (y-axis).

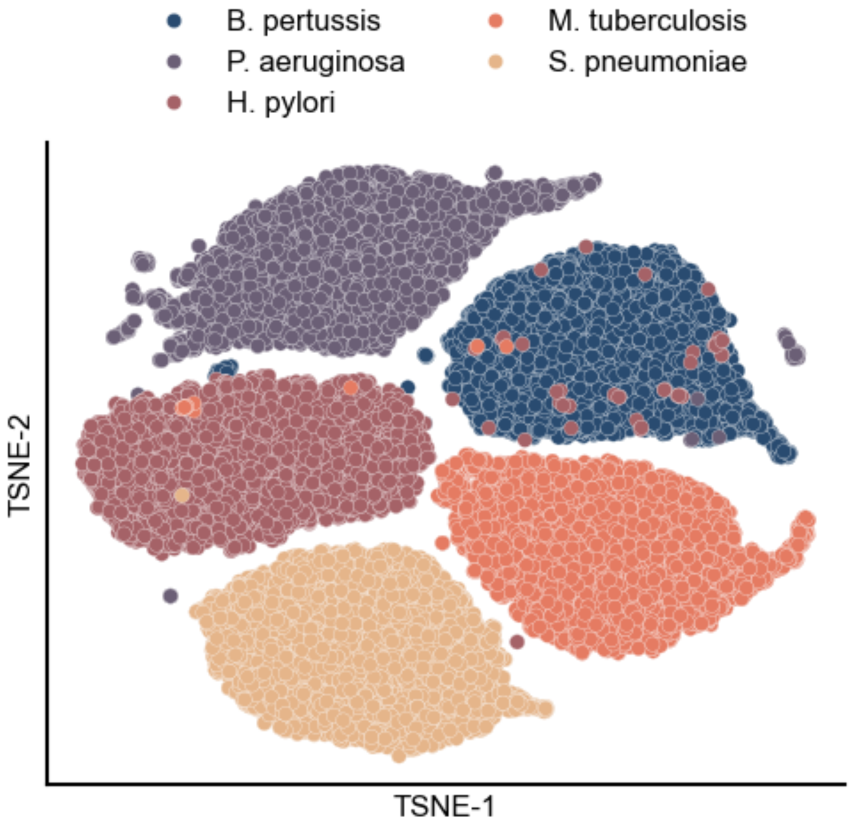


FIG. S2. **Variational autoencoder compression of five bacterial pathogens.** OptGPSampler was used to obtain $N = 5000$ samples ($k = 100$) from five bacterial GEMs selected from Plata et al, Nature, 2015. A variational autoencoder (VAE) was trained on the common $N = 494$ reactions shared across all species. The final embedding dimension $D$ =8 was reached via 5 linearly decreasing ($D = 494$, $D = 397$, $D = 300$, $D = 203$, $D = 107$, $D = 8$) fully connected layers and the VAE was trained with PyTorch for 100 epochs with default hyperparameters using the Adam optimizer; t-SNE embeddings were employed for 2-dimensional visualization. The embeddings captures the separation of metabolic spaces across species. While the learned representation displays a small number of outliers, overall we found that all 5 pathogens retain distinct clustering despite the VAE only being trained on non species-specific reactions.

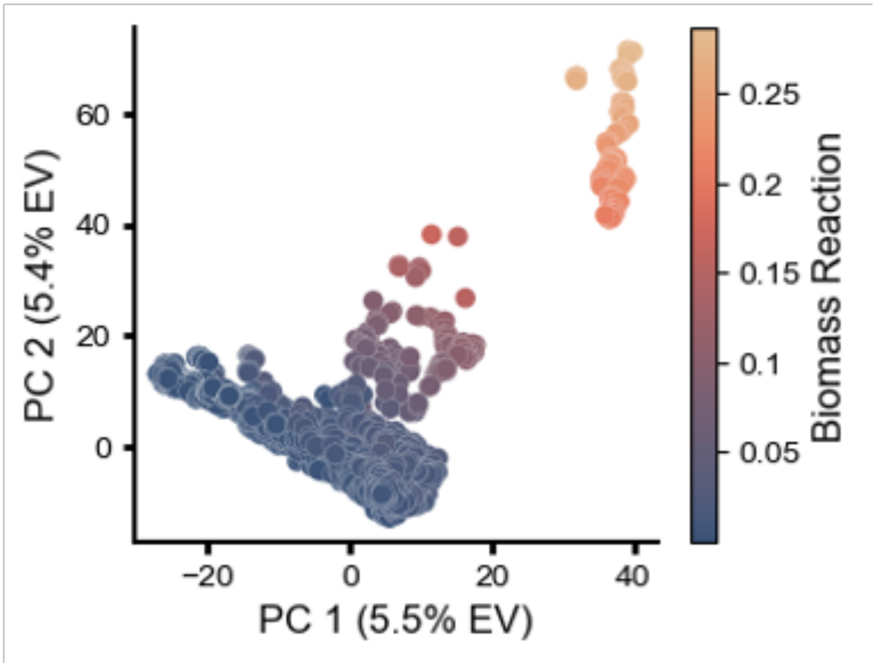


FIG. S3. **PCA representation of flux cone of wild type *E. coli* metabolism.** The unmodified (WT) iML1515 model was sampled 5000 times with a step size k=100.The resulting data was normalized using zero mean and unit variance and Principal Component Analysis (PCA) was performed. 621 PCs were required to explain 95% of variance in the scaled data. The first two PCs accounted for 5.5% and 5.4% of variance, respectively. The biomass reactions were removed from the PC calculations (see Supplementary Table S3) but samples on the plot are colored based on the value of the forward biomass reaction (BIOMASS_Ec_iML1515_core_75p37M).

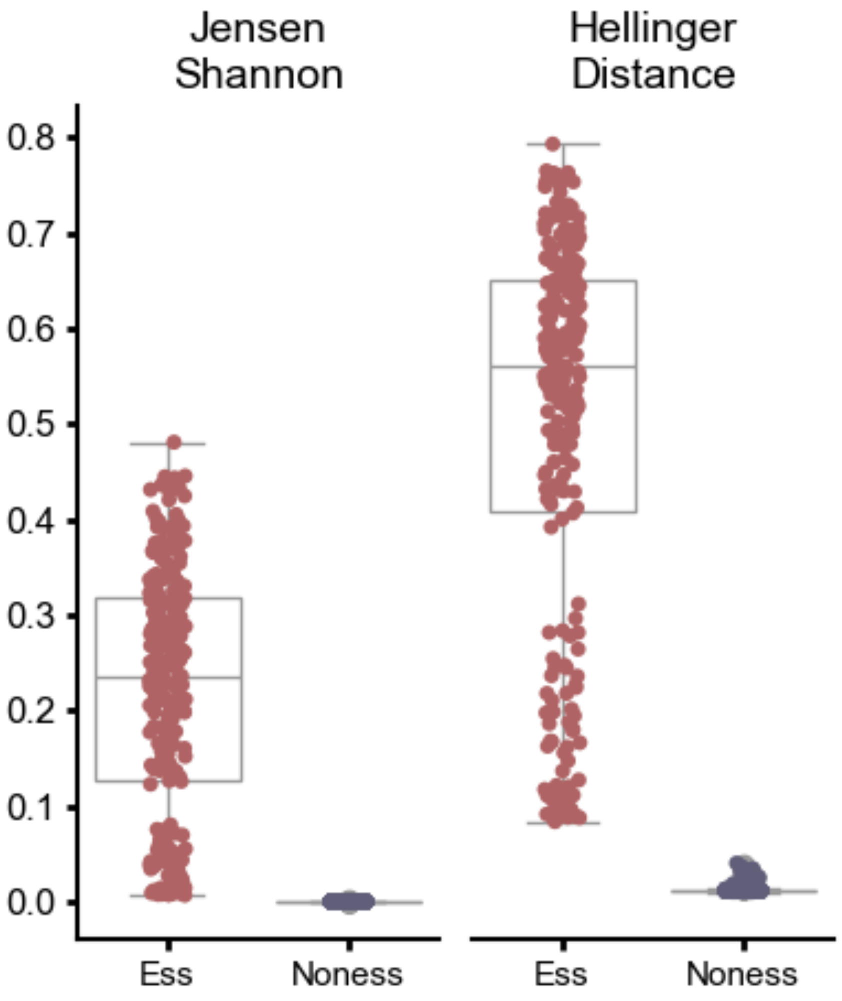


FIG. S4. **Complete classifier sample scoring.** We sought to exploit the excellent predictive performance of FCL to define a distance metric between deletions and the wild type strain. To this end, we retrained FCL on all *E. coli* gene deletions ($N$ =1,502 genes), and computed the distribution of prediction scores for all flux samples in the wild type and each deletion strain. We then queried the model with flux samples of the wild-type GEM and each deletion cone (100 samples/cone), to produce distributions of prediction scores for the WT and each deletion strain. We scored each strain with the Jensen-Shannon divergence and Hellinger distance between the score distributions of each deletion and the wild type. We found statistically significant differences in score distribution of non-essential and essential genes, which reinforces the conclusion that perturbations to the flux cone are indicative of gene essentiality (one-sided t-test with p 0.05).

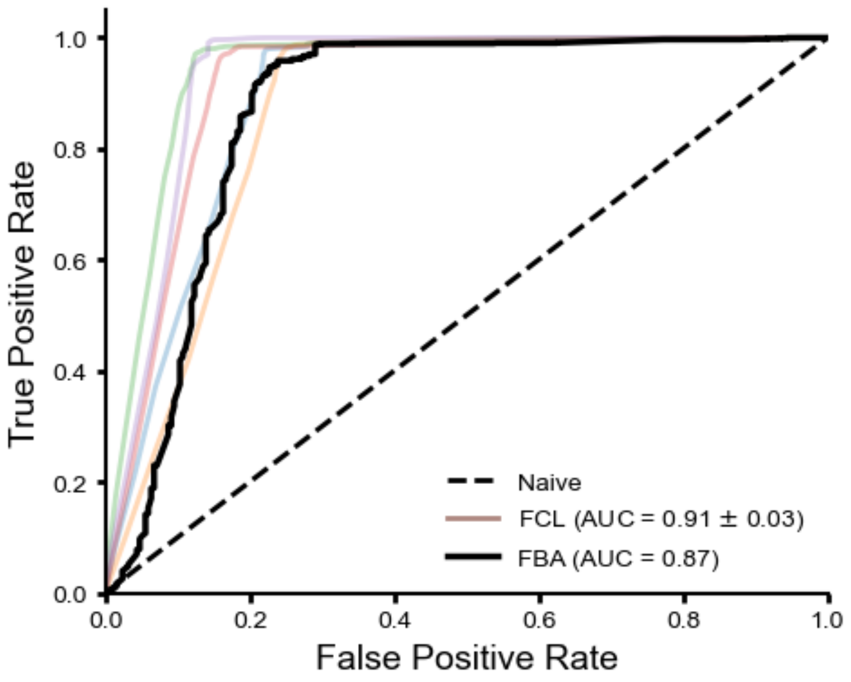


FIG. S5. **Receiver operating characteristic (ROC) curves for *E. coli* essentiality prediction.** FCL predictions were computed on a held-out set of test genes (20%) across five training repeats for a single model (100 samples/cone density) selected randomly from those in Figure 2A–B; FBA predictions were computed for all genes.

| GEM | Number of genes | Number of reactions | Stoichiometric nullity |
|---|---|---|---|
| iML1515 | 1516 | 2712 | 867 |
| iJO1366 | 1367 | 2583 | 817 |
| iAF1260 | 1261 | 2382 | 752 |
| ijr904 | 904 | 1075 | 332 |
| Yeast9 | 1162 | 4130 | 1539 |
| iCHO2291 | 2291 | 6236 | 2390 |

TABLE S1. Summary of genome-scale metabolic models employed in this study.

| GEM | Sampled KOs | Total S | Density per cone | Step size | Subsampling | Size of NPZ file |
|---|---|---|---|---|---|---|
| iML1515 | 1502 | 150313 | 101 | 100 | 50 | 3.06Gb |
| iJO1366 | 1318 | 163680 | 124 | 5000 | 1 | 3.17Gb |
| iAF1260 | 1214 | 150784 | 124 | 5000 | 1 | 2.69Gb |
| ijr904 | 866 | 107508 | 124 | 5000 | 1 | 0.89Gb |
| Yeast9 | 1121 | 143716 | 124 | 1 | 1 | 4.43Gb |
| iCHO2291 | 2290 | 291028 | 127 | 1 | 1 | 13.53Gb |

TABLE S2. Summary of model sampling details. All models were sampled with the same step size, except iML1515 which employed a smaller step size to generate more samples and the subsampled for the analysis in Figure 2.

| GEM | Biomass Reactions Removed [IDs] |
|---|---|
| iML1515 | BIOMASS_Ec_iML1515_core_75p37M [2669], BIOMASS_Ec_iML1515_core_75p37M_reverse_35685 [2670] |
| iJO1366 | BIOMASS_Ec_iJO1366_core_53p95M [19], BIOMASS_Ec_iJO1366_core_53p95M_reverse_5c8b1 [14] |
| iAF1260 | BIOMASS_Ec_iAF1260_core_59p81M [926] |
| ijr904 | BIOMASS_Ecoli [269], BIOMASS_Ecoli_reverse_bf7a1 [270] |
| Total Common Reactions | 864 |

TABLE S3. Biomass reactions removed for *E. coli* models and their common reactions.

| Task | Class imbalance | Class breakdown |
|---|---|---|
| *E. coli* essentiality | 83/17 | N=1252 non-essential, N=251 essential |
| Yeast essentiality | 86/14 | N=964 non-essential, N=157 essential |
| CHO essentiality | 83/17 | N=1898 non-essential, N=392 essential |
| Yeast production | 17/67/16 | N=138 low, N=545 medium, N=128 high |

TABLE S4. Class imbalances for each task.

| Organism | Model | Accuracy | Precision | Recall | F1 Score | PR AUC | ROC AUC |
|---|---|---|---|---|---|---|---|
| Yeast | FBA | 0.81 | 0.91 | 0.97 | 0.89 | 0.92 | 0.69 |
| Yeast | FCL (5-fold CV) | 0.90±0.008 | 0.91±0.008 | 0.98±0.010 | 0.94±0.007 | 0.94±0.021 | 0.78±0.065 |
| Yeast | FCL (test) | 0.89 | 0.89 | 0.99 | 0.94 | 0.93 | 0.73 |
| CHO | FBA | 0.85 | 0.86 | 0.97 | 0.91 | 0.86 | 0.62 |
| CHO | FCL (5-fold CV) | 0.85±0.014 | 0.86±0.010 | 0.97±0.008 | 0.91±0.008 | 0.91±0.029 | 0.72±0.068 |
| CHO | FCL (test) | 0.84 | 0.86 | 0.97 | 0.91 | 0.86 | 0.65 |

TABLE S5. Classification performance metrics for Flux Cone Learning (FCL) and Flux Balance Analysis (FBA) in yeast (*S. cerevisiae*) and Chinese Hamster Ovary (CHO) cells (Figure 3).

| Model Name | Baseline | Resampled | Balanced | Both | Max % improvement |
|---|---|---|---|---|---|
| HistGradientBoostingClassifier | 11.4 | 13.6 | 14.1 | 14.6 | 28.3 |
| LinearSVC | 23.8 | 24.0 | 27.2 | 26.8 | 27.2 |
| LogisticRegression | 23.3 | 26.2 | 29.5 | 28.9 | 14.2 |
| RandomForestClassifier | 18.1 | 11.7 | 19.1 | 18.4 | 5.5 |

TABLE S6. Accuracy for high producer deletion yeast deletion strains for each class balancing methods in Figure 4. Models were assessed across all genes in a held-out, class-stratified 20% test set ($N$ =649 deletions).
